# Organic soil amendment with Spent Mushroom Substrate results in fungal colonisation, alters bacterial-fungal co-occurrence patterns and improves plant productivity

**DOI:** 10.1101/788935

**Authors:** Fabiana S. Paula, Enrico Tatti, Camilla Thorn, Florence Abram, Jude Wilson, Vincent O’Flaherty

## Abstract

In agricultural systems based on organic fertilisers, the activity of prokaryotes and fungi is essential for degradation of complex substrates and release of nutrients for plant uptake. Understanding the dynamics of microbial communities in these systems is, therefore, desirable for designing successful management strategies aiming to optimise nutrient availability and to improve plant productivity. Of particular interest is how the microbial inoculum provided by an organic substrate persists in the soil and interacts with soil and plant microbiomes, as these processes may affect the long-term benefits of organic amendments. We aimed to investigate how these dynamics occurred in soil treated with stabilised spent mushroom substrate (SMS), a soil amendment rich in nutrients and complex organic matter. We carried out a 14 weeks soil trial to assess the plant growth promoting properties of the SMS and to monitor the successional processes of the resulting SMS-soil communities compared to a mineral amended control. Bacterial and fungal communities were analysed by high-throughput sequencing at both DNA and RNA (cDNA) levels. Using a combination of computational tools, including SourceTracker and Network analysis, we assessed the persistence of SMS-derived taxa in soil, and the changes in co-occurrence patterns and microbial community structure over time. Prokaryotic and fungal communities presented remarkably distinct trajectories following SMS treatment. The soil prokaryotic communities displayed higher levels of resilience to the changes introduced by SMS treatment and rapidly tended toward a soil-like profile, with low persistence of SMS-derived prokaryotes. In contrast, the SMS fungal community had greater success in soil colonisation during the time monitored. SMS treatment promoted an increase in the participation of fungi in the highly connected fraction of the active community, including fungal taxa of SMS origin. We observed the presence of highly connected microbial guilds, composed by fungal and bacterial taxa with reported capabilities of complex organic matter degradation. Many of these taxa were also significantly correlated with the organic matter content and plant yield, suggesting that these highly connected taxa may play key roles not only in the community structure, but also in the plant-soil system under organic fertilisation.

## Introduction

Organic residues from agriculture and the food industry provide sustainable alternatives to synthetic fertilisers and thus promote the circular economy (Grimm and Wösten, 2018). Residues from the mushroom industry, herein referred to as spent mushroom substrate (SMS), are currently treated as a waste, despite having great potential for use in the agricultural sector due to high organic matter, nitrogen (N), phosphorus (P), potassium (K) contents, as well as large scale availability (Jordan et al., 2008; Roy et al., 2015). A recent study demonstrated that a stabilised SMS product was able to support plant growth and significantly improved grass yield in soil-free horticultural mixes (Paula et al., 2017). In addition, the benefits of using unprocessed SMS as an organic fertiliser and soil conditioner have already been documented (Courtney and Mullen, 2008; Hackett, 2015). However, the mechanisms underpinning the effects of this organic amendment on plant performance and soil microbiology have yet to be explored in depth.

In contrast to mineral fertilisers, when organic residues are applied to soil a large portion of the nutrients are present as complex molecules, not available for direct plant uptake. The decomposition of the organic matter (OM) is, therefore, a key step to releasing nutrients from the biomass. The first stages of this process are carried out by fungi and bacteria capable of secreting hydrolytic enzymes, which break down complex carbohydrates (Berlemont, 2017). Thereafter, the course of the microbial community succession affects OM transformation (Fontaine et al., 2003) and, therefore, impacts nutrient availability (Hellequin et al., 2018).

An increasing number of studies aimed to disentangle the complexity of microbial communities in soils under fertilisation systems (Bonanomi et al., 2016; Paul Chowdhury et al., 2019; Randall et al., 2019), and how it may affect plant health. Yet community assembly and dynamics are still far from being predictable. Particularly in organic agriculture, responses of the soil microbiome to biotic and abiotic factors introduced by organic amendments can vary considerably.

In addition to nutrients and physical structure, organic amendments may also provide a microbial inoculum to the soil. How these microbes interact with the soil community, persist in the environment and affect key soil functions is still very uncertain. Factors such as the resilience of the soil microbiome, i.e. the rate it responds to changes (Martiny et al., 2017), may affect the persistence of the inoculum. Such knowledge is essential to exploit the full potential of organic systems (Agler et al., 2016) and to design microbiome management strategies aiming to improve soil health, plant productivity and resistance to environmental stresses. For instance, fungi and bacteria have been proposed to play different roles in decomposition processes, mainly as a result of divergent metabolic capabilities, substrate preference (reviewed by de Menezes et al., 2017) and growth rate/mode (de Graaff et al., 2010). The complexity of these interactions is likely further increased in a system containing microbial inputs from different sources, such as soil and organic amendments. Here we used SMS as a soil amendment and organic fertiliser and evaluated its plant growth promotion properties, as well as its effects on the rhizosphere microbial communities. To better understand the microbial dynamics, we aimed to test whether fungi and prokaryotes from the SMS would persist in the soil, and if so, how they might integrate with the resident soil microbiota and potentially affect soil community structure. To tackle these questions, we followed the community succession at both DNA and RNA levels, and used a combination of computational tools, including SourceTracker (Knights et al., 2011) and network analysis (Faust and Raes, 2012) to evaluate, respectively, the persistence of SMS microbes and the interactions of the newly formed community. We searched for highly connected taxa and consortia with the assumption that these microbes may play an important role in the community structure (Busby et al., 2017), with a focus on groups potentially involved in the degradation of complex organic matter and nutrient release. Taxa with high connectivity have been suggested to mediate the effects of abiotic factors on plant microbiomes (Faust and Raes, 2012). Finally, we explored how the relative abundance of these hub taxa correlated with soil parameters and plant yield.

## Methods

### Experimental setup and soil sampling

Italian ryegrass (*Lolium multiflorum*) was grown in top soil of a silt loam texture and pH 7.0. Stabilised SMS (SMS) (Paula et al., 2017) was used as an organic soil amendment in the absence of additional nutrient sources. Two organic treatments, low (OL; 60 g/L) and high (OH; 110 g/L) were applied, to represent two frequently used application rates, 45 and 85 tons/ha, respectively. A mineral treatment (M) contained N-0.33; P-0.04; K-0.32 in g/L, aiming to supply a similar P content as the OH treatment, without exceeding P application limits (European Union Regulations, 2014), and also maintaining the recommended NPK ratio. In the untreated control (U), no source of NPK was provided. The soil mixes were homogenised thoroughly, placed into 1 L pots and sown with 40 mg of ryegrass seeds. Each treatment was replicated in 25 pots. The pots were organised in a completely randomised plot design in a greenhouse facility (Radharc, Galway, Ireland), and after each sampling time, pots were re-randomised. Automatic irrigation was employed to keep soil moisture between 60 and 80 %. The trial was conducted from March to June 2015, with average daily temperature 14.9 ± 1.5 (mean ± SD). The NPK contents in the SMS are presented in **Table S1**. Additional nutrient characterisation of the SMS substrate has been published previously (Paula et al., 2017). The physical and chemical properties of the soil during the trial are presented in **Table S2**.

Grass yield was measured using four herbage harvests taken from ten (non-destructive) replicates per treatment at weeks 5, 8, 11 and 14 after fertiliser application. Grass was cut 1 cm above the substrate level and dried at 55°C for 72 hours to determine plant dry weight per pot. For soil analysis, three destructive replicates were collected per sampling time. The plant dry weight of the destructive replicates was also determined to be used in correlation analysis with microbial parameters. Soil microbial community analyses were carried out on soil collected at time zero and at weeks 8 and 14. Plants were carefully removed from pots and gently shaken to remove loose soil from intact roots. Thereafter, soil firmly attached to the roots was removed to obtain the rhizosphere fraction. Samples were immediately frozen in liquid nitrogen to preserve RNA and were stored at −80°C until processing.

### *Analytical method*s

Available P and K were quantified by standard procedures (CEN - EN 13650). Mineral nitrogen was measured in 2 M KCl extract (1:5 w/v) where NH_4_^+^ was determined according to (Kandeler and Gerber, 1988), and NO_2_^-^ and NO_3_^-^ were quantified as described previously (Shand et al., 2008; Keeney and Nelson, 1982). Air dried samples were used to determine pH and electrical conductivity (EC) in 1:5 soil: water dilution (w/v). Samples were oven dried at 105°C to assess dry matter content. Dry material was used to measure Kjeldahl nitrogen (N) (Lang, 1958) and total organic matter (TOM; Bremner and Mulvaney, 1982).

### Nucleic acid extraction and processing

DNA and RNA were co-extracted from 0.5 g of soil using the method described by Griffiths et al. (2000), modified with the addition of casein to increase the nucleic acid yield, as demonstrated by Wang et al. (2012). To remove remaining PCR inhibitors, the OneStep PCR inhibitor removal kit (Zymo Research, Irvine, CA, USA) was employed. Nucleic acids from two extractions per replicate were combined and eluted in 40 µl of PCR grade water. Quality was verified on a 1% agarose gel and using a Nanodrop 2000 spectrophotometer (Thermofisher Scientific, Waltham, USA) set for determining absorbance at the following wavelengths: 230, 260, 280 and 320 nm. An aliquot of the purified nucleic acid was treated with DNase (Turbo DNA-free kit; Thermo Fisher Scientific, Waltham, MA, USA) and RNA was reverse transcribed to cDNA using SuperScript III Reverse Transcriptase Kit (Invitrogen, Carlsbad, California, United States). Nucleic acid concentrations were assessed using Qubit dsDNA HS Assay and Qubit RNA HS Assay Kits (Thermo Fisher Scientific, Waltham, MA, USA).

### Taxonomic profiling of the microbial communities

For taxonomic profiling of the soil fungal communities, the Internal Transcribed Spacer 2 (ITS2) region was amplified with the region-specific primers ITS3F/ITS4R (White et al., 1990). To investigate the prokaryotic communities, the primers 515F/806R (Caporaso et al., 2011) were used to amplify the V4 region of the 16S rRNA gene. Paired-end sequencing was performed on an Illumina MiSeq (Illumina, Inc. San Diego, California) 2×300 flow cell, at Research and Testing Laboratories (Lubbock, TX), following standardised procedures. ITS and 16SrRNA sequence data were deposited on NCBI’s Sequence Read Archive under the accession number SUB3044384: http://www.ncbi.nlm.nih.gov/biosample/7345359.

Sequence processing was conducted in Qiime platform (Caporaso et al., 2010). For 16S rRNA fragment analyses, merged forward and reverse reads were quality-filtered using *multiple_split_libraries_fastq.py* with quality threshold of phred 19. Putative chimeric sequences were identified using the *de novo* method in USEARCH (Edgar, 2010; Edgar et al., 2011). USEARCH was also used for open reference OTU clustering, with the RDP consensus taxonomy assigner, employing SILVA (release 132 - 10.04.2018) as a reference database. ITS sequences were treated according to the optimised pipeline described by Taylor et al. (2016), with the following steps differing from the 16SrRNA analyses: forward and reverse reads were not joined and analyses were carried out using forward read data only – as a consequence of the low quality of the reverse reads; following quality filtering, the ITS2 fragment was extracted using ITSx (Bengtsson-Palme et al., 2013); OTU clustering was performed with open reference method providing the UNITE database (ver7_dynamic_12.01.2017) as the reference; taxonomy was assigned with the blast method, and non-fungi sequences were excluded from the dataset. The prokaryotic DNA and RNA datasets had totals of 1,570,832 and 1,029,199 reads, after quality control, respectively, while for fungi, the DNA and RNA datasets had 1,935,165 and 3,143,854 reads, after quality control, respectively. Singleton sequences were removed and within each of the four datasets (16S rRNA/ITS; DNA/ RNA) the number of sequences were rarefied to equalise the sampling depth per sample in an effort to reduce sequencing biases.

### Quantitative PCR

The number of 16S rRNA gene and ITS2 fragment copies were assessed in soil DNA samples using qPCR with the region-specific primer sets used for taxonomic profiling. The reaction mixtures for both fragments consisted of 0.6 μL of each primer (10 uM), 10 μL of 2 x SsoFast EvaGreen Supermix, 4 ng of template DNA and water in a total volume of 20 μL. Copy number quantification was performed using a LightCycler 480 Instrument (Roche, Penzberg, Germany). The temperature program for 16S rRNA fragments consisted of 98 °C for 2 min, followed by 40 cycles of denaturation at 98 °C for 5 s and annealing/elongation at 60 °C for 20 s. As for ITS2, initial 95 °C for 2 min, followed by 40 cycles of 95 °C for 30 s, 55 °C for 30 s and 72 °C for 60 s. Standard curves for 16S rRNA and ITS2 fragments were generated using serial dilutions of *Pseudomonas aeruginosa* and *Penicillium brevicompactum* DNA, respectively, from 10^2^ to 10^8^ copies/μL.

### Data analysis

Statistical analyses were performed in R version 3.4.4 (R core team, 2018), unless otherwise stated. Differences in grass yield at each sampling time were assessed using one-way ANOVA and Tukey’s multiple comparisons tests. Two-way ANOVA was used to test the effects of treatment and sampling time on gene copy number and on Shannon diversity index. The effect of treatment and time on the microbial community composition was tested using a PERMANOVA model (Anderson, 2001).

SourceTracker version 1.0.1 (Knights et al., 2011) was used to partition the different sources that explain the composition of the microbial communities in SMS treated soils across time. This method uses abundance data to seek for low or moderate source environment endemicity. It also assigns as ‘unknown’ grouping when part of a sink sample is unlike any of the provided sources or when it is unable to identify discriminatory taxonomic signatures among sources (Henry et al., 2016; Knights et al., 2011).

To search for patterns of co-occurrence among microbial taxa, we used Co-occurrence Network Inference (CoNet) (Faust and Raes, 2012). Associations among OTUs and higher-level taxa were calculated using the Pearson, Spearman, Kendall, Bray Curtis and Kullback-Leibler correlation methods simultaneously (Wang et al., 2017). The analysis proceeded with the top 1000 positive interactions. The significance of the edges was assessed by permutation and the p-values were corrected for multiple testing with Benjamini-Hochberg procedure (Benjamini and Hochberg, 1995), and edges with p-values below 0.05 were retained. Interactions supported by a minimum of 2 methods were considered true. Edge and node files were input to Gephi (Bastian et al., 2009) version 0.9.2 to construct the final network. Spearman’s correlations of hub taxa with environmental factors were assessed using Hmisc package (Harrel and Dupont, 2016) and the results were plotted using corrplot (Wei and Simko, 2017).

## Results and discussion

### SMS as a soil organic amendment

Stabilised SMS was applied to soil as an organic amendment in the absence of additional fertilisers. We previously showed that SMS was able to support plant growth and increase plant yield in a soil-free horticultural mix (Paula et al., 2017). Here we assessed the ability of SMS to support *Lolium multiflorum* growth in a soil trial, and further investigated the effects of this organic amendment on soil biotic and abiotic characteristics. Results were compared with mineral amended and untreated soils over the 14-week trial.

A rapid increase in grass yield was seen only in the mineral (M) amended soils at the initial harvests (weeks 5 and 8), which peaked at week 8 (**Figure 1**). At the third harvest (week 11), there was an increase in grass yield in the organic-high (OH) treatment, which was not statistically different from M. At the fourth and final harvest (week 14) yields from OH were statistically significantly higher than from M. Organic-low (OL) yields also improved as the trial proceeded, but at a smaller scale than OH, indicating a dose dependent response of the treatment.

**Figure 1.**
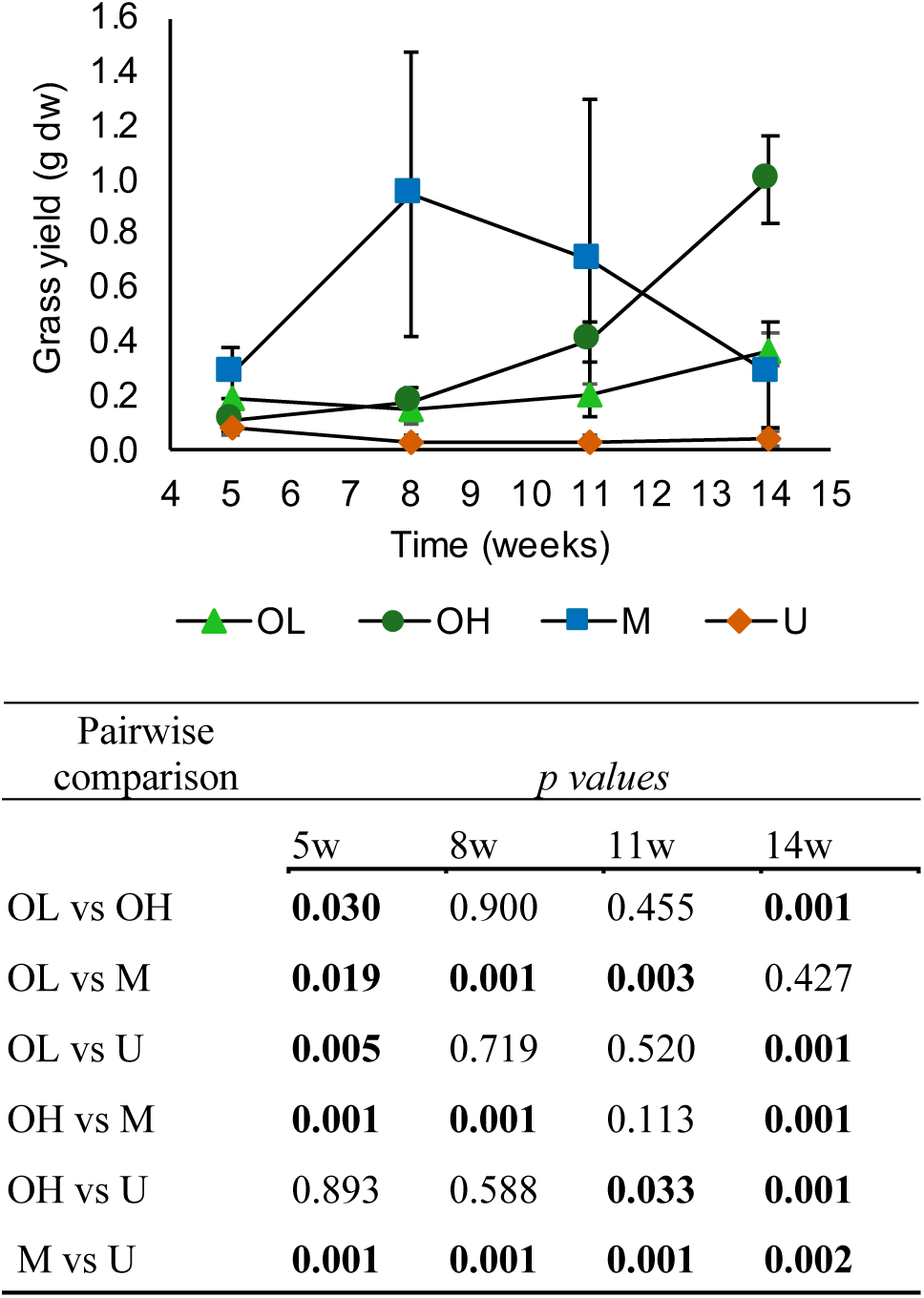
Changes in grass yield in response to organic and mineral treatments. Grass dry weight was measured after 5, 8, 11 and 14 weeks of trial. The values represent the mean ± SD of 10 replicates per treatment. OL, Organic-Low; OH, Organic-High; M, Mineral; U, Untreated. The table presents the p values of the pairwise comparisons at each time point (ANOVA and Tukey’s post hoc test). p<0.05 are marked in bold.

The rapid response observed in M was expected, due to the high levels of plant available NPK, and in particular nitrogen (**Table S2**). Despite the short-term advantages of mineral fertilisers, large scale applications of these products are often associated with contamination of groundwater or surface waters, as the available nutrients may be leached from soil before they can be taken up by plants. Organic agriculture, in contrast, is known to promote long-term productivity owing to the slow release of nutrients (Castaldi et al., 2004). In fact, incorporation of SMS to soil caused an increase not only in the available NPK stocks, but also in total N and organic matter (**Table S2**). These nutrient sources require microbial activity in order to release the mineral fractions for plant uptake. Simultaneously, the addition of complex organic matter may also cause an increase in soil organic matter degradation, as a result of a priming effect (Bernard et al., 2007).

The changes in plant yield presented above for organic and mineral treatments during this 14-week trial are a representation of field profiles that can be observed in long term agriculture systems. It is therefore an interesting model to investigate dynamics of soil microbial communities in response to organic and mineral fertilisation, and to hone our knowledge on how these dynamics may contribute to the plant-soil system.

### Dynamics of the rhizosphere microbial communities in response to SMS amendment

To investigate changes in the grass rhizosphere microbial communities in response to SMS amendment, we assessed the diversity of prokaryotes and fungi by sequencing 16S rRNA and ITS genes, respectively. In addition, the potentially active community was examined by sequencing the transcripts for those regions, allowing us to target the fraction of the communities likely responding to environmental changes. While studying rRNA and ITS transcripts, there are some known limitations to be acknowledged, including the presence of ribosomal RNA reported in dormant cells, and variability in concentration in active cells across taxa (Blazewicz et al., 2013). However, distinct responses (Meyer et al., 2019) and higher correlations with environmental variations are often reported for rRNA, in comparison to rDNA (Zhang et al., 2014). This is particularly critical in a short-term trial with microbial inputs from two different sources, here SMS and soil. In this study we did not attempt to make quantitative comparisons between the communities assessed by DNA or RNA approaches. Instead, we integrated both datasets with the assumption that the active community is likely to be enriched in the RNA fraction, and that combining DNA and RNA data may offer more robust information on the dynamics and establishment of the new SMS-soil mixed communities.

Irrespective of the sample type or treatment analysed, the prokaryotic communities were largely dominated by Bacteria, with Archaea representing less than 1%. Archaeal relative abundance was below 0.1% for the SMS substrate and all soils at time zero, and tended to increase with time in all treatments, with dominance of the phylum Nanoarchaeota (**Figure S1**). Members of this poorly characterised phylum, known to contain small genomes and to allegedly living in symbiosis with other organisms, have been detected in a variety of environments (Munson-McGee et al., 2015).

At the phylum level, several differences were observed among the samples regarding their bacterial composition. Untreated soil prokaryotic communities were dominated by Proteobacteria in both DNA (51%) and RNA (60%) fractions, followed by Bacteroidetes, Verrucomicrobia and Acidobacteria (**Figure S1**). This is in agreement with previous reports showing that Proteobacteria is often found to be dominant in rhizosphere communities (reviewed by Philippot et al., 2013). In the SMS substrate community, Proteobacteria was still the dominant phylum, with relative abundances of 27% (DNA) and 39% (RNA), while large fractions of the communities were also assigned to Bacteroidetes (DNA-25%; RNA – 17%) and Actinobacteria (DNA – 18%; RNA – 14%). Differences between the DNA and RNA datasets were more evident at lower taxonomical ranks: in the initial soil communities, the most abundant class in the DNA data was Gammaproteobacteria (26%), while at the RNA level Deltaproteobacteria was dominant with a mean relative abundance of 43%. At time zero, the SMS treated soils (OH and OL) presented a mixed soil-SMS community profile, which tended to be more similar to untreated soils towards the end of the trial for both DNA and RNA (**Figure S1**).

Ascomycota was the dominant fungal phylum across all samples, particularly in the SMS where it accounted for above 99% of the OTUs (**Figure S2**). The dominance of Ascomycota in the SMS was also observed in the initial communities of OH and OL. Organically amended soils (OH and OL) followed different successional trajectories, when compared to U and M treated soils. The relative abundance of Basidiomycota tended to increase with time in all treatments, particularly in the active fraction, but it was higher in M and U, compared to OL and OH, throughout the trial, although previous studies have correlated their abundance with increase in organic matter (Peay et al., 2017). By contrast, a significant increase of the saprophytic Chytridiomycota relative abundance was only observed in OH and OL at week 14 (**Figure S2**). This effect, however, was not mirrored in RNA data.

The community similarities were further analysed by NMDS ordination and PERMANOVA analysis. Soil prokaryotic and fungal communities were significantly affected by SMS treatment at both DNA and RNA levels (**Figure 2**; **Table S3**). The changes observed are likely to be a combined result of bioaugmentation and stimulation of specific microbial groups by the organic matter input (Hellequin et al., 2018). By contrast, the mineral treatment did not cause significant changes in fungal or prokaryotic community composition relative to the untreated soils. Multivariate analysis revealed that the treatment effect was stronger in fungal communities, while changes in prokaryotic communities presented higher variation with time (**Table S3**). This is consistent with the high turnover rates usually presented by bacterial communities, in contrast with the slower growth modes of fungi (de Graaff et al., 2010). We also assessed how the community within each treatment changed with time and observed that the prokaryotic communities from the organic treatments presented the highest variation, while the mineral treatment appeared to be less affected in all datasets **(Table S3**).

**Figure 2.**
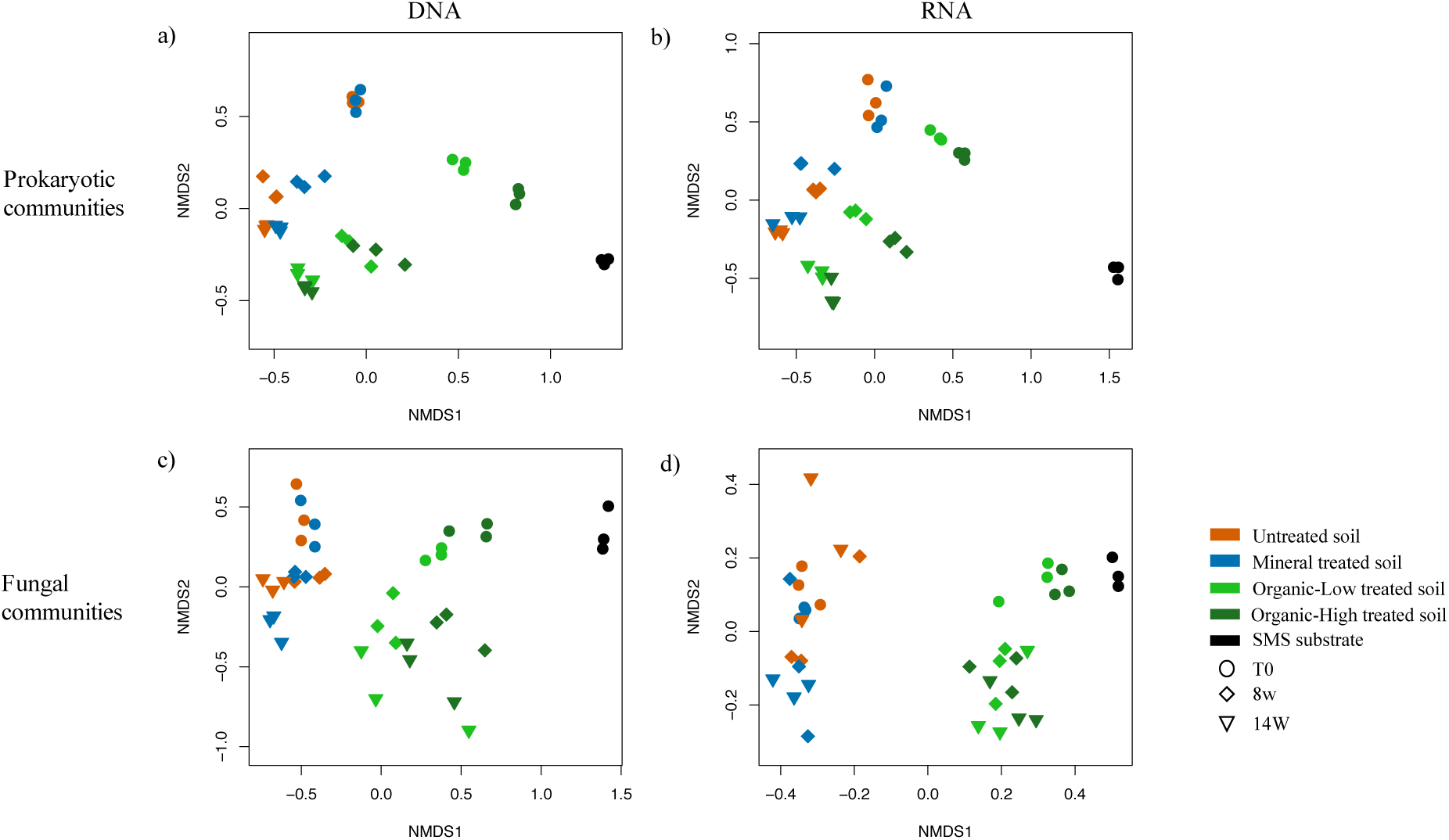
Nonmetric multidimensional scaling of prokaryotic (a, b) and fungal (c, d) community composition, at DNA (a, c) and RNA levels (b, d). Dissimilarity index: Bray-Curtis; Stress: a) 0.042; b) 0.053; c) 0.091; d) 0.095. t0, time zero; 8w, 8 weeks; 14w, 14 weeks.

When changes in microbial community diversity were investigated as a function of time or treatment, we observed a similar response: prokaryotic diversity varied more as a function of time than treatment, while the opposite was observed for fungal diversity (**Figure S3; Table S4**). The diversity of prokaryotes was initially reduced in response to the SMS treatment, compared to U and M, but values in all treatments converged with time. Although different profiles were observed for the potentially active community at time zero, these also tended to equalise towards the end of the experiment. The diversity of soil fungi, initially reduced with SMS amendment, also showed recovery during the trial, but remained at significantly lower levels in the SMS treated soils, at the DNA level. Similar trends were observed for RNA, yet without statistical significance. Fungal diversity indices have previously been shown to be more negatively impacted than bacterial diversity during the process of straw decomposition (Banerjee et al., 2016). The lower microbial diversity observed in SMS treated soils at time zero is likely a result of an increase in dominant groups provided by SMS. The effects of the SMS community and nutrient inputs over the indigenous soil microbiota are observed in the early stages of the trial, as has been reported for other types of organic amendments (Hellequin et al., 2018). In this 100 day-experiment it was possible to observe a return of the prokaryotic community diversity to untreated soil-like levels, which did not occur in previously reported shorter term experiments using organic amendments (Hellequin et al., 2018). The fungal community, in contrast, may need a longer time to recover diversity. Alternatively, SMS-borne fungi may be favoured by the recalcitrant resources available, thus sustaining their dominance.

The total abundance of 16S rRNA and ITS fragments was assessed by qPCR. As expected, amendment with SMS caused an initial increase in both prokaryotic and fungal abundance, compared to the untreated soil (**Figure 3 a, b**), owing to bioaugmentation with the microbial pools present in the SMS (**Figure 3 d**). As the trial proceeded, a rise in total fungal count occurred in the SMS treated soils, while U and M did not present significant changes. No such effect was observed for the abundance of prokaryotes (**Figure 3 a, b; Table S5**). The fungi to prokaryote ratio evolved differently under the different treatments as the trial progressed. There was a reduction in the fungal proportions in M treated soils, in agreement with previous research demonstrating that fertilisers promote bacterial-dominated communities (Beare et al., 1997; de Vries et al., 2006). Despite the sustained bacterial dominance across all treatments, OH communities tended to become enriched with fungi (**Figure 3c**). A longer trial would be required to evaluate whether or not fungi become the dominant microbes in SMS treated soil, as observed for other organic amendments (Banerjee et al., 2016). Fungi dominated soils are proposed to increase carbon sequestration due to slower turnover, higher C-use efficiency and the presence of more chemically recalcitrant components in fungal biomass (Strickland and Rousk, 2010). Therefore, as well as its crop nutritive value, SMS could represent a sustainable means of improving carbon sequestration (Favoino and Hogg, 2008).

**Figure 3.**
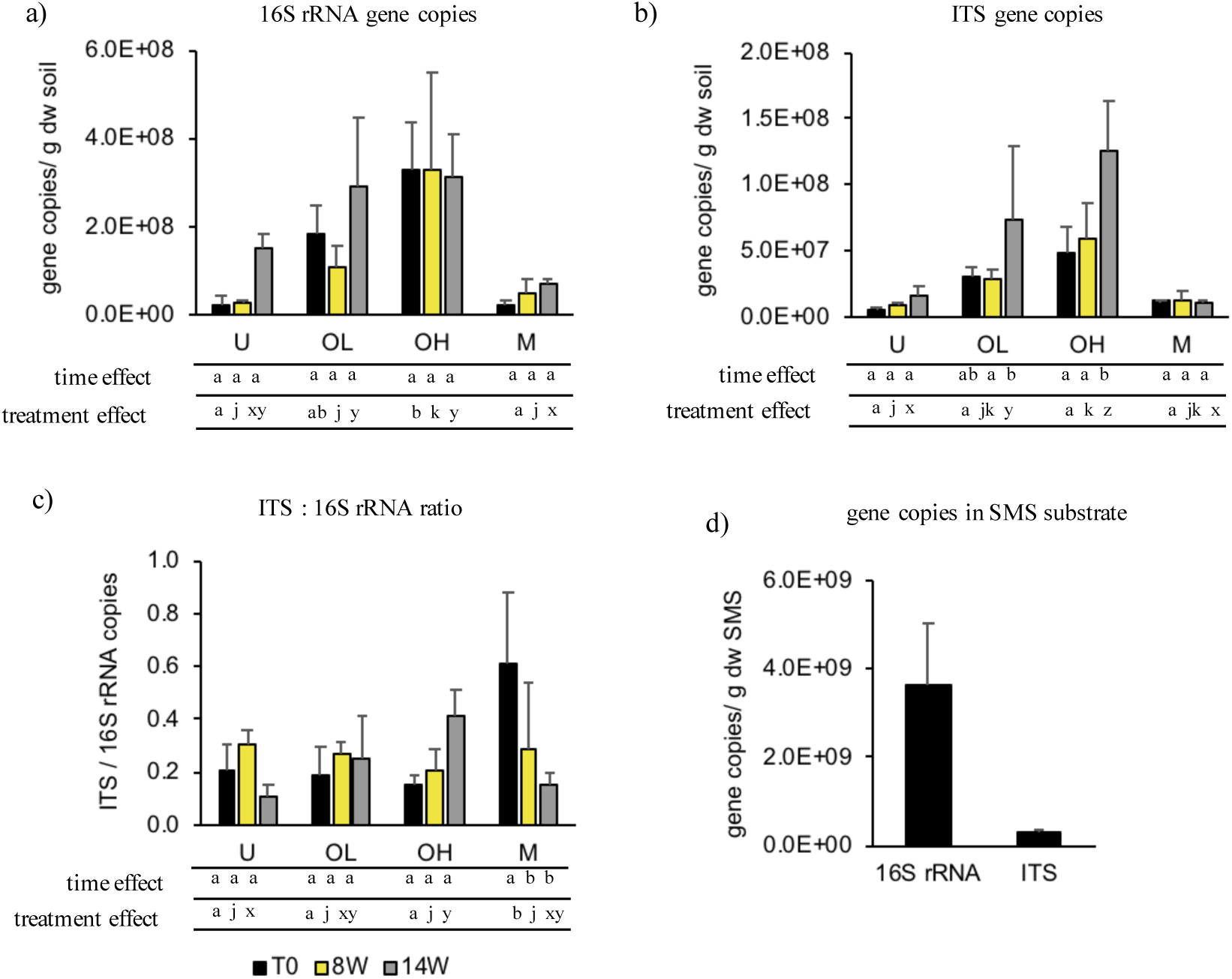
Prokaryote (a) and Fungi (b) abundance in soils under the different treatments, assessed by qPCR. c) fungi: prokaryotes ratio; d) Prokaryote and Fungi abundance in the SMS substrate. Statistics: Two-way ANOVA is presented in Table S6. Tukey’s post hoc test is presented on the bottom of figures a-c. Time effect (within same treatment): time points not sharing the same letter (a,b,c) were significantly different from each other; Treatment effect (at same time point): treatments not sharing the same letter (a,b,c / jkl / xyz) were significantly different from each other.

### Persistence of SMS microbial communities in soil

The results of community composition (**Figure 2**), diversity (**Figure S3**) and abundance (**Figure 3**) indicated different paths for prokaryotic and fungal community succession. Despite being subjected to different treatments, the prokaryotic diversity in all soils tended to converge to a similar status and their community succession also seemed to follow comparable trajectories. These observations suggest that the prokaryotic pool added by bioaugmentation may not have persisted in the soil. Simultaneously, there were indications that the soil prokaryotic community presented resilience to the disturbance introduced by SMS amendment, similarly to previous reports evaluating other organic treatments (Lourenço et al., 2018). By contrast, the effects of the SMS fungi inoculum seem to be more persistent over time. We asked to what extent these contrasting patterns could be observed in the community composition, and used SourceTracker (Knights et al., 2011) to estimate the proportions of SMS treated soil communities throughout the trial that were attributed to SMS in soil samples. This approach allowed us to investigate the persistence of SMS fungi and prokaryotes in the soil environment and to explore the soil microbiome resilience under these conditions. SourceTracker allows microbial communities to be examined in terms of the potential sources of the microbiota. Here we used SMS and untreated soil from T0 as sources contributing organisms to the SMS-treated soils, referred to as sinks.

It should be noted that the microbial biomass between these two sources varied considerably, where gene copy numbers (per gram dry weight) were two orders of magnitude more abundant in SMS than in the soil (**Figure 3d**). Consequently, at time zero, high proportions of the community composition in OH were attributed to SMS source, explaining over 60 and 80% of the prokaryotic (**Figure 4a**) and fungal (**Figure 4c**) communities, respectively. Interestingly, when the communities were analysed at the RNA level, SMS-associated fractions contributed to approximately 40% of prokaryotic communities (**Figure 4b**) and above 90% of fungal communities (**Figure 4d**) at time zero. The fate of these SMS and soil derived proportions was different between prokaryotic and fungal groups. With progression of the trial, prokaryotes attributed to SMS reduced considerably in relative abundance, with a concomitant increase in soil-derived prokaryotes. In contrast, SMS fungi were more persistent and presented only a small reduction with time, at both DNA and RNA levels, suggesting a successful establishment in the soil (**Figure 4**). A similar trend was observed for OL treatment (**Figure S4**).

**Figure 4.**
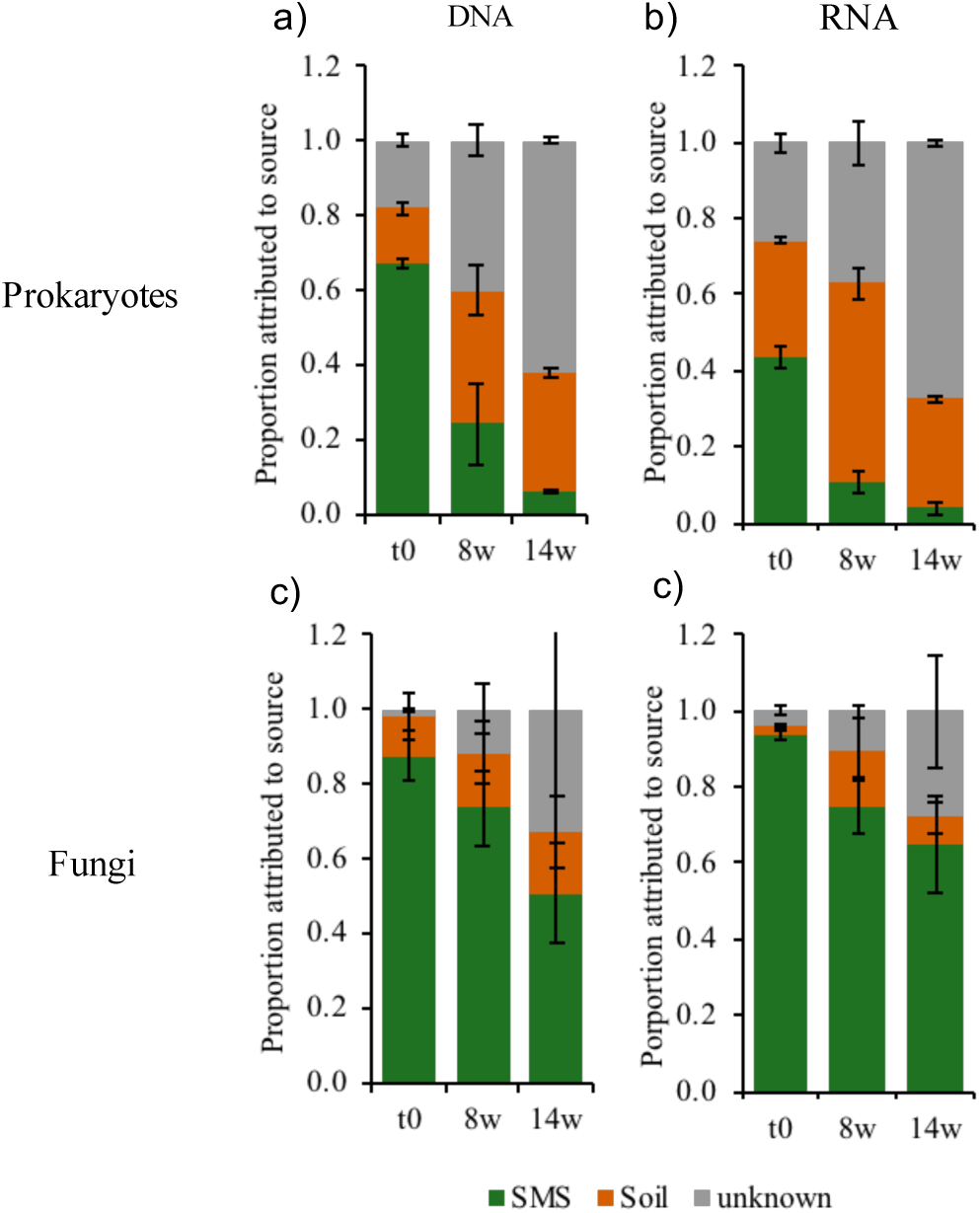
Source proportions of prokaryotic (a, b) and fungal (c, d) communities estimated for OH soils at time 0, 8 weeks and 14 weeks, using SourceTracker. Values represent mean proportions ± SD. a & c – DNA; b & d – RNA.

Finally, the community fraction attributed to unknown sources, i.e. undefined origin, increased with time for both communities, but to a lesser degree for fungi. Microbiota identified by SourceTracker as unknown in source refers to those groups which the tool cannot attribute to any of the sources provided or is unable to discriminate signatures among sources (Henry et al., 2016; Knights et al., 2011). We hypothesise that these microbes could be present in our sources, but in such low numbers that they are either not detected by the sequencing depth employed herein, or they are represented by singletons, which are removed as common practice in analysing sequencing data. In other words, they might represent transiently very rare taxa, present either in the SMS or the soil, and possibly in a dormant state (Aanderud et al., 2015). Changes in season, time, temperature, moisture (Shade et al., 2014) and notably growth of the ryegrass with the associated root exudates may have allowed for them to rise to detectable levels. Also inputs of nutrients from the SMS may have provided further resource availability promoting increases in their relative abundances.

Bacteria and fungi can produce hydrolytic enzymes to deconstruct complex carbohydrates (Berlemont, 2017). Whole genome investigations have explored the phylogenetic distribution of potential degraders. There is a high diversity of bacteria classified as opportunists, which produce only β-glucosidases that perform the last step of polysaccharide degradation, and therefore, process smaller substrates released by other taxa (Berlemont and Martiny, 2013). Bacteria equipped with the complete set of enzymes for full breakdown of the polysaccharides, including cellulases and β-glucosidases are less common. Some phyla with higher frequencies of this phenotype are Actinobacteria, Bacteroidetes, Firmicutes and Proteobacteria (Berlemont and Martiny, 2015). SMS-treated soils presented significantly higher relative abundance of Actinobacteria and Firmicutes, when compared to untreated soils (**Figures S5**). Within the Actinobacteria phylum, the family Streptosporangiaceae presented a remarkably higher relative abundance in OH at time zero (**Figure S6a)**. Despite the higher values in OH and OL until week 14, the relative abundance of this group declined sharply with time, in comparison to U and M. A similar trend was observed for the Firmicutes families Bacillaceae (**Figure S6b)** and Paenibacillaceae (**Figure S6c)** at the DNA level.

Fungi, in contrast, are regarded as generalists and genome inspections have indicated that the genotypes required to fully degrade multiple complex polysaccharides, such as cellulose, xylan and chitin, are phylogenetically widespread (Berlemont, 2017). In addition, the role of fungi in the degradation of recalcitrant organic matter is often highlighted due to their ability to breakdown lignin, which is rarely reported for bacteria (Strickland and Rousk, 2010). Fungal groups with recognised cellulose degrading capabilities were among the dominant taxa in the SMS substrate. From those, the family Chaetomiaceae is often linked to the degradation of complex organic matter in soils and have been identified as one of the main groups responsible for the difference between untreated and organic-treated soils (Banerjee et al., 2016). In fact, this family was dominant in SMS treated soils and tended to increase or sustain their high relative abundance with time, when compared to U and M (**Figure S7**). More remarkable, however, was the high relative abundance of this group in the RNA data, which was above 50% at all sampling times. The high relative abundance of Chaetomiaceae in the potentially active community may suggest a possible role of this group in the organic matter turnover and nutrient release in SMS treated soils.

### Co-occurrence patterns of the microbial community in response to SMS treatment

To further investigate the dynamics of fungi and prokaryotes in response to SMS treatment, we used network analysis to visualise co-occurrence patterns among taxa. Prokaryotic and fungal data were integrated to explore their potential interactions and roles in the structure of the communities under different treatments. While a number of studies have aimed to decipher bacterial co-occurrence dynamics in the environment (Ling et al., 2016; Wang et al., 2017), the roles of fungi in microbial networks are still poorly explored, as only a few surveys have evaluated the co-occurrence of fungal taxa either separately or combined with other organisms (Agler et al., 2016; Banerjee et al., 2019). Here we not only investigated the effects of the organic treatment over the soil cross-domain community networks, but also, we explored how these dynamics occurred at both DNA and RNA levels. Finally, we used the network approach to explore how microbes of SMS origin integrated into the soil community.

Each network was built using data from rhizosphere samples collected at weeks 8 and 14, to encompass temporal variability within treatments. It has been reported that the importance of certain microbial taxa in the community structure may change with spatiotemporal variability (Banerjee et al., 2018). Therefore, investigations of co-occurrence patterns across time may reveal profiles that persist in the environment, and hence are likely to have an important role in microbiome functioning.

The network analysis revealed remarkably distinct profiles for total and potentially active communities (**Figure 5**). The networks built from RNA datasets showed consistently higher number of nodes, edges, average degree and average path length, indicating higher interaction complexity than DNA networks (**Table S6**). In contrast, DNA networks tended to present higher modularity and average clustering coefficient, which are parameters related to the tendency to form local clusters (Banerjee et al., 2019; Zhou et al., 2011). In addition, in the RNA data, highly connected taxa, also called “microbial hubs”, were more frequent. The assessment of RNA-derived networks possibly enabled a visualisation of the fast dynamics and turnover of the communities.

**Figure 5.**
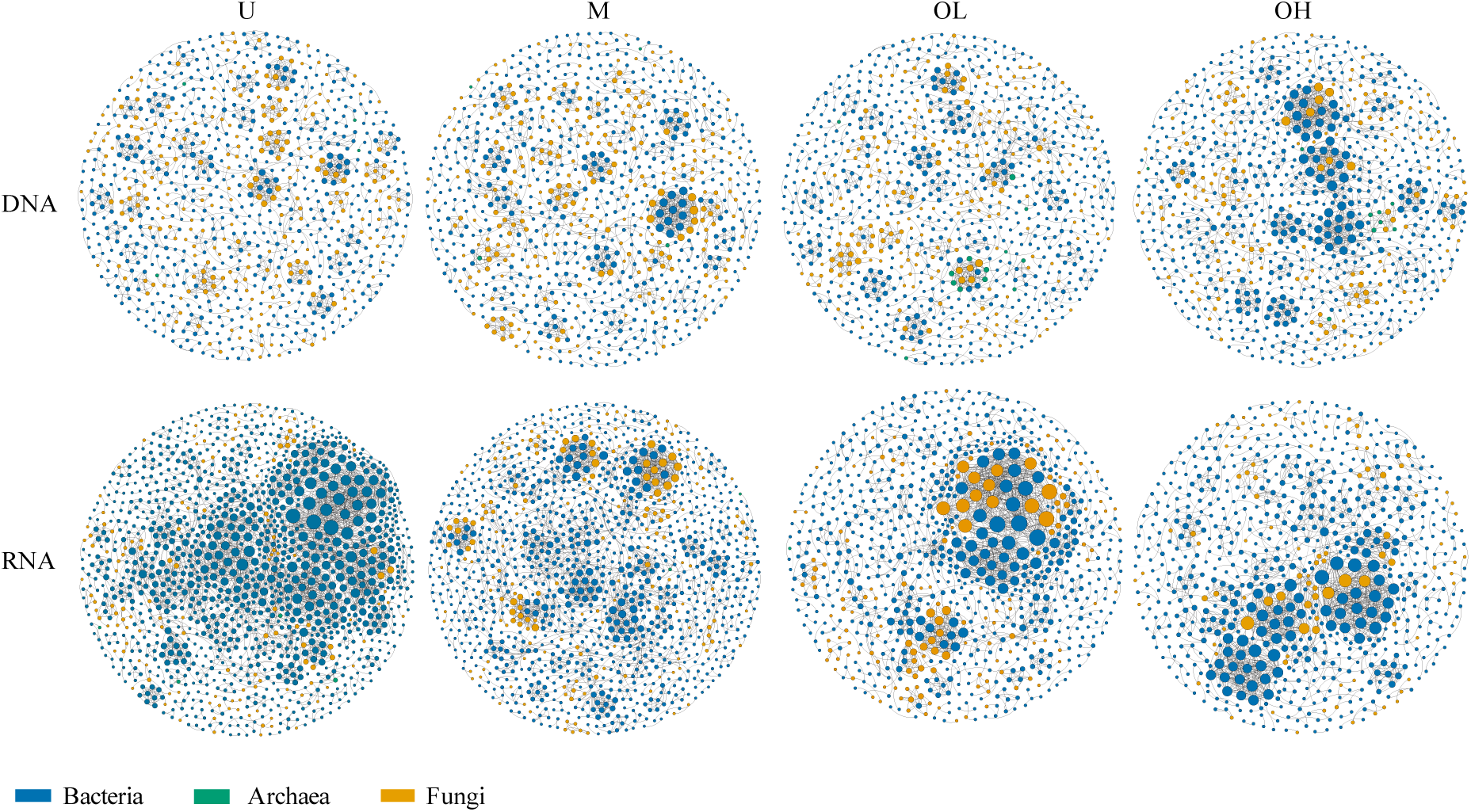
Rhizosphere microbial community co-occurrence patterns. Nodes (circles) represent operational taxonomic units (OTU) or higher taxa levels and edges (lines) represent significant co-occurrence between two nodes. Node scale defined by connectivity degree. Blue, Bacteria; Orange, Fungi; Green, Archaea. U, Untreated; OL, M, Mineral; Organic-Low; OH, Organic-High.

The effects of the treatments on network topology were found to be different between DNA and RNA (**Figure 5**). At the DNA level, M, U and OL networks contained mostly poorly connected taxa (i.e. low degree nodes), while a few hub nodes (herein defined as nodes with > 20 edges) were observed in OH. At the RNA level, the untreated soil network presented numerous high-degree OTUs, 34 of which were classified as hubs (**Table S7**). Interestingly, all those nodes belonged to prokaryote taxa. RNA networks from OL and OH contained 39 and 37 hub taxa, respectively. But unlike untreated soils, these comprised both bacteria and fungi (**Table S7**). The SMS treatment, therefore, seems to have promoted an increase in the participation of fungi in the highly connected fraction of the active community, as well as in their connectivity with prokaryotes. The latter point was further demonstrated by the assortativity index, here used to assess whether taxa are more likely to interact with taxonomically related groups (Kurtz et al., 2015). When the assortativity index was calculated for the RNA networks to compare interaction at the Domain level, the lowest value observed for the treatment OH: 0.14, indicating that taxa therein had more interactions outside their own taxonomic Domain. The index values for the other treatments, OL, M and U were 0.24, 0.30 and 0.27, respectively.

To understand how soil and SMS microbes interacted in the newly formed communities of SMS treated soils, we further explored the network of their potentially active taxa (**Figure 5**). Among the fungal hub nodes, we observed the presence of several Chaetomiaceae OTUs (**Table S7**), some of which were detected in the SMS substrate, but not in the soil initial community (**Figure S8**), suggesting that these groups may have originated from the SMS inoculum. Therefore, not only can SMS-borne fungal taxa persist and thrive in the soil **(Figure 4)**, but some groups also seem to become part of the highly connected fraction of the potentially active community. In contrast, none of the bacterial hubs were detected in the SMS substrate community, suggesting they were part of the indigenous soil microbiota. Soil-borne bacterial hub taxa tended to increase their relative abundance with time (**Figure S8**), and included taxa known for containing cellulose-degrading strains, such as the Proteobacteria families Bulkholderiaceae (Wilhelm et al., 2019) and Polyangiaceae (Garcia and Müller, 2014). Hub microbes with ecologically relevant roles in the community are more likely to function as keystones (Agler et al., 2016), which are taxa that may exert major influences on community structure and functioning and ecosystem processes (Agler et al., 2016; Banerjee et al., 2018). Several studies have identified keystone species in soil microbiomes linked to organic matter decomposition (Banerjee et al., 2016) and nutrient transformation (Li et al., 2017). In SMS treated soils, we observed fungal and bacterial hubs that were assigned to taxa known to comprise groups with reported capabilities for complex organic matter degradation. These guilds with highly connected and functionally redundant taxa might have a higher impact on broad processes, such as organic matter decomposition (Banerjee et al., 2018).

In the OL network, two OTUs assigned to the arbuscular mycorrhizal fungi (AMF) family Claroideoglomeraceae were also among the highly connected taxa. AMF are symbionts that can provide multiple benefits to the plant by facilitating nutrient transfer, increasing tolerance to stressors and improving soil quality and structure (Varela-Cervero et al., 2016). The role of AMF as keystone taxa has been suggested in soils under different agricultural systems (Banerjee et al., 2019). Other hub microbes detected in this study belong to taxa previously identified as keystone taxa in rhizospheres and other soil microbiomes., including Rhizobiales, Verrucomicrobia, Bacteroidetes and Acidobacteria (Banerjee et al., 2018).

Hub microbes are suggested to influence the whole microbiome by both direct and indirect mechanisms. Keystone taxa may affect the abundance and activity of other groups, thereby impacting community assembly (Agler et al., 2016), structure and performance (Banerjee et al., 2018). Alternatively, in plant microbiomes, hub microbes have been shown to mediate the interaction between the microbiome and abiotic factors (Agler et al., 2016). Investigating how rhizosphere hub taxa interact with environmental factors may contribute to elucidating the complexity of the plant-soil system and to devising plant microbiome management strategies. We investigated how the relative abundance of hub microbes (degree >20) correlated with changes in environmental factors across OH and M samples (**Figure 6**). Several hub microbes, from both treatments, positively correlated with NO_3_^-^. Interestingly, N and P presented only a few significant correlations with hub taxa, all of which were negative. Organic matter content presented positive correlations mainly with hub taxa from the OH treatment, including all fungal taxa. Grass yield correlated positively with ITS:16S rRNA ratio, suggesting that the fungal biomass may play a crucial role in plant productivity in this system. Only hub taxa from OH correlated with grass yield. Interestingly, many of these taxa (OTU) were also positively correlated with OM content, comprising groups of both soil and SMS origin (**Figure S8**). Banerjee et al. (2016) reported significant associations of bacterial and fungal keystone taxa with OM decomposition. The data presented here do not provide any mechanistic link between taxa abundance/activity, organic matter transformation and plant productivity. However, the presence of several decomposers as hub taxa, in addition to their correlation with organic matter content and plant yield, may suggest a potential role of these microbes in the plant-soil system under SMS treatment.

**Figure 6.**
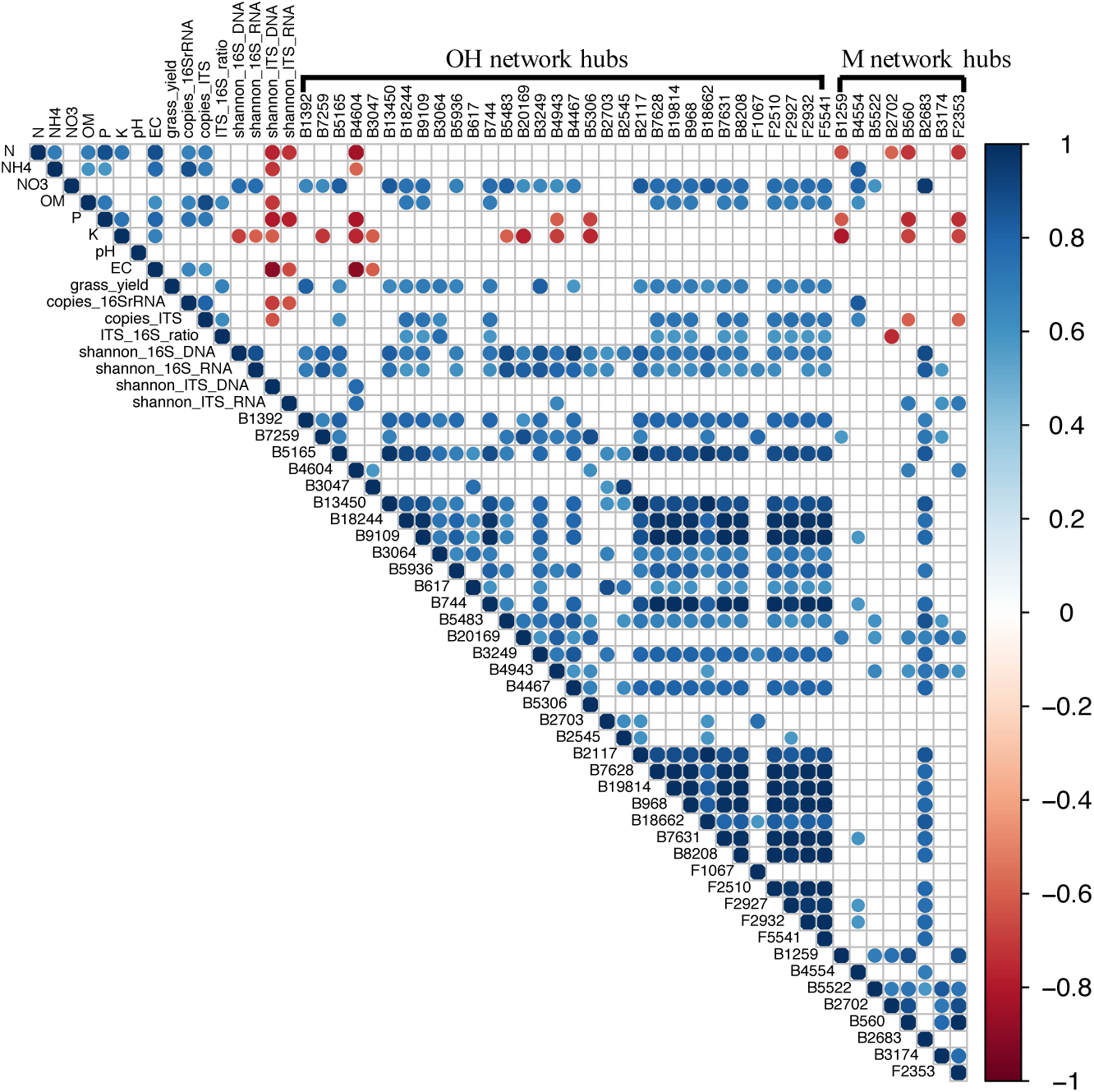
Spearman correlations among community parameters and environmental factors. OTUs classified as hub taxa (degree >20) in OH and M treatments were tested for their correlation with microbial and environmental factors across these two treatments. “B” and “F” hubs refer to bacterial and fungal OTUs respectively. Significant correlations (p<0.05) are marked in blue (positive) and red (negative). For taxonomic classification of the hub taxa, see Table S7.

The results presented above demonstrated that SMS amendment successfully supported plant growth. Besides the nutrients and physical properties provided by this substrate, the microbial inocula may have contributed to the benefits observed. In addition, bacterial and fungal communities followed distinct dynamics in soil under organic amendment with SMS substrate. The data suggested that the soil prokaryotic community had higher levels of resilience following SMS treatment and rapidly tended toward a soil-like profile, while the fungi community provided by the SMS seemed to have greater success in soil colonisation. Some of the SMS fungi may have the potential to perform important roles in the soil community structure, as they were among the highly connected microbes. Together with soil-derived microbes, these fungi formed hub guilds, with the potential to affect community structure. Finally, some of these hub microbes also seemed to correlate with plant yield and organic matter content. These findings could provide an initial framework for soil microbiome management strategies aiming to explore persistence of microbial inocula, resilience of soil communities and microbial groups with potential impacts on the soil community structure and function.

## Acknowledgments

The authors thank Dr. Edna Curley for contributing with initial discussion, and the members of the Compost Analysis Laboratory at Monaghan Mushrooms, for analytical support.

## Author Contributions

F.S.P, E.T. and J.W wrote the grant proposal; F.S.P, E.T., F.A., J.W. and V.O.F. designed the research; F.S.P, E.T. and C.T performed the experiments; F.S.P analysed the data and wrote the manuscript. All authors contributed with discussion and critically reviewed the manuscript.

## References

Aanderud, Z.T., Jones, S.E., Fierer, N., Lennon, J.T., 2015. Resuscitation of the rare biosphere contributes to pulses of ecosystem activity. Frontiers in Microbiology 6, 24. doi:10.3389/fmicb.2015.00024

Agler, M.T., Ruhe, J., Kroll, S., Morhenn, C., Kim, S.-T., Weigel, D., Kemen, E.M., 2016. Microbial Hub Taxa Link Host and Abiotic Factors to Plant Microbiome Variation. PLOS Biology 14, e1002352. doi:10.1371/journal.pbio.1002352

Anderson, M.J., 2001. A new method for non-parametric multivariate analysis of variance. Austral Ecology 26, 32–46. doi:10.1111/j.1442-9993.2001.01070.pp.x

Banerjee, S., Kirkby, C.A., Schmutter, D., Bissett, A., Kirkegaard, J.A., Richardson, A.E., 2016. Network analysis reveals functional redundancy and keystone taxa amongst bacterial and fungal communities during organic matter decomposition in an arable soil. Soil Biology and Biochemistry 97, 188–198. doi:10.1016/J.SOILBIO.2016.03.017

Banerjee, S., Schlaeppi, K., van der Heijden, M.G.A., 2018. Keystone taxa as drivers of microbiome structure and functioning. Nature Reviews Microbiology 16, 567–576. doi:10.1038/s41579-018-0024-1

Banerjee, S., Walder, F., Büchi, L., Meyer, M., Held, A.Y., Gattinger, A., Keller, T., Charles, R., van der Heijden, M.G.A., 2019. Agricultural intensification reduces microbial network complexity and the abundance of keystone taxa in roots. The ISME Journal 13, 1722–1736. doi:10.1038/s41396-019-0383-2

Bastian, M., Heymann, S., Jacomy, M., n.d. Gephi: An Open Source Software for Exploring and Manipulating Networks Visualization and Exploration of Large Graphs.

Beare, M.H., Hu, S., Coleman, D.C., Hendrix, P.F., 1997. Influences of mycelial fungi on soil aggregation and organic matter storage in conventional and no-tillage soils. Applied Soil Ecology 5, 211–219. doi:10.1016/S0929-1393(96)00142-4

Bengtsson-Palme, J., Ryberg, M., Hartmann, M., Branco, S., Wang, Z., Godhe, A., De Wit, P., Sánchez-García, M., Ebersberger, I., de Sousa, F., Amend, A.S., Jumpponen, A., Unterseher, M., Kristiansson, E., Abarenkov, K., Bertrand, Y.J.K., Sanli, K., Eriksson, K.M., Vik, U., Veldre, V., Nilsson, R.H., 2013. Improved software detection and extraction of ITS1 and ITS2 from ribosomal ITS sequences of fungi and other eukaryotes for analysis of environmental sequencing data. Methods in Ecology and Evolution 4, /a-n/a. doi:10.1111/2041-210X.12073

Benjamini, Y., Hochberg, Y., 1995. Controlling the False Discovery Rate: A Practical and Powerful Approach to Multiple Testing, Source: Journal of the Royal Statistical Society. Series B (Methodological).

Berlemont, R., 2017. Distribution and diversity of enzymes for polysaccharide degradation in fungi. Scientific Reports 7, 222. doi:10.1038/s41598-017-00258-w

Berlemont, R., Martiny, A.C., 2015. Genomic Potential for Polysaccharide Deconstruction in Bacteria. Applied and Environmental Microbiology 81, 1513–1519. doi:10.1128/AEM.03718-14

Berlemont, R., Martiny, A.C., 2013. Phylogenetic distribution of potential cellulases in bacteria. Applied and Environmental Microbiology 79, 1545–54. doi:10.1128/AEM.03305-12

Bernard, L., Mougel, C., Maron, P.-A., Nowak, V., Lévêque, J., Henault, C., Haichar, F. el Z., Berge, O., Marol, C., Balesdent, J., Gibiat, F., Lemanceau, P., Ranjard, L., 2007. Dynamics and identification of soil microbial populations actively assimilating carbon from ^13^ C-labelled wheat residue as estimated by DNA- and RNA-SIP techniques. Environmental Microbiology 9, 752–764. doi:10.1111/j.1462-2920.2006.01197.x

Blazewicz, S.J., Barnard, R.L., Daly, R.A., Firestone, M.K., 2013. Evaluating rRNA as an indicator of microbial activity in environmental communities: limitations and uses. The ISME Journal 7, 2061–2068. doi:10.1038/ismej.2013.102

Bonanomi, G., De Filippis, F., Cesarano, G., La Storia, A., Ercolini, D., Scala, F., 2016. Organic farming induces changes in soil microbiota that affect agro-ecosystem functions. Soil Biology and Biochemistry 103, 327–336. doi:10.1016/J.SOILBIO.2016.09.005

Bremner, J.M. and Mulvaney, C.S., 1982. Nitrogen-Total. In: Methods of soil analysis. Part 2. Chemical and microbiological properties, Page, A.L., Miller, R.H. and Keeney, D.R. Eds., American Society of Agronomy, Soil Science Society of America, Madison, Wisconsin, 595-624.

BS EN 13650:2001-Soil improvers and growing media - Extraction of Aqua Regia soluble elements. British Standard, London

Busby, P.E., Soman, C., Wagner, M.R., Friesen, M.L., Kremer, J., Bennett, A., Morsy, M., Eisen, J.A., Leach, J.E., Dangl, J.L., 2017. Research priorities for harnessing plant microbiomes in sustainable agriculture. PLOS Biology 15, e2001793. doi:10.1371/journal.pbio.2001793

Caporaso, J.G., Kuczynski, J., Stombaugh, J., Bittinger, K., Bushman, F.D., Costello, E.K., Fierer, N., Peña, A.G., Goodrich, J.K., Gordon, J.I., Huttley, G.A., Kelley, S.T., Knights, D., Koenig, J.E., Ley, R.E., Lozupone, C.A., McDonald, D., Muegge, B.D., Pirrung, M., Reeder, J., Sevinsky, J.R., Turnbaugh, P.J., Walters, W.A., Widmann, J., Yatsunenko, T., Zaneveld, J., Knight, R., 2010. QIIME allows analysis of high-throughput community sequencing data. Nature Methods 7, 335–6. doi:10.1038/nmeth.f.303

Caporaso, J.G., Lauber, C.L., Walters, W.A., Berg-Lyons, D., Lozupone, C.A., Turnbaugh, P.J., Fierer, N., Knight, R., 2011. Global patterns of 16S rRNA diversity at a depth of millions of sequences per sample. Proceedings of the National Academy of Sciences of the United States of America 108 Suppl 1, 4516–22. doi:10.1073/pnas.1000080107

Castaldi, P., Garau, G., Melis, P., 2004. Influence of compost from sea weeds on heavy metal dynamics in the soil-plant system. Fresenius Environmental Bulletin 13 (11).

Courtney, R.G., Mullen, G.J., 2008. Soil quality and barley growth as influenced by the land application of two compost types. Bioresource Technology 99, 2913–2918. doi:10.1016/J.BIORTECH.2007.06.034

de Graaff, M.-A., Classen, A.T., Castro, H.F., Schadt, C.W., 2010. Labile soil carbon inputs mediate the soil microbial community composition and plant residue decomposition rates. New Phytologist 188, 1055–1064. doi:10.1111/j.1469-8137.2010.03427.x

de Menezes, A.B., Richardson, A.E., Thrall, P.H., 2017. Linking fungal–bacterial co-occurrences to soil ecosystem function. Current Opinion in Microbiology 37, 135–141. doi:10.1016/J.MIB.2017.06.006

de Vries, F.T., Hoffland, E., van Eekeren, N., Brussaard, L., Bloem, J., 2006. Fungal/bacterial ratios in grasslands with contrasting nitrogen management. Soil Biology and Biochemistry 38, 2092–2103. doi:10.1016/J.SOILBIO.2006.01.008

Edgar, R.C., 2010. Search and clustering orders of magnitude faster than BLAST. Bioinformatics 26, 2460–2461. doi:10.1093/bioinformatics/btq461

Edgar, R.C., Haas, B.J., Clemente, J.C., Quince, C., Knight, R., 2011. UCHIME improves sensitivity and speed of chimera detection. Bioinformatics 27, 2194–2200. doi:10.1093/bioinformatics/btr381

European Union (Good Agricultural Practice for Protection of Waters) Regulations 2014.

Faust, K., Raes, J., 2012. Microbial interactions: from networks to models. Nature Reviews Microbiology 10, 538–550. doi:10.1038/nrmicro2832

Favoino, E., Hogg, D., 2008. The potential role of compost in reducing greenhouse gases. Waste Management & Research 26, 61–69. doi:10.1177/0734242X08088584

Fontaine, S., Mariotti, A., Abbadie, L., 2003. The priming effect of organic matter: a question of microbial competition? Soil Biology and Biochemistry 35, 837–843. doi:10.1016/S0038-0717(03)00123-8

Garcia, R., Müller, R., 2014. The Family Polyangiaceae, in: The Prokaryotes. Springer Berlin Heidelberg, Berlin, Heidelberg, pp. 247–279. doi:10.1007/978-3-642-39044-9_308

Griffiths, R.I., Whiteley, A.S., O’Donnell, A.G., Bailey, M.J., Bernillon, D., LeGall, F., Jeannin, P., Nesme, X., Simonet, P., 2000. Rapid method for coextraction of DNA and RNA from natural environments for analysis of ribosomal DNA- and rRNA-based microbial community composition. Applied and Environmental Microbiology 66, 5488–91. doi:10.1128/aem.66.12.5488-5491.2000

Grimm, D., Wösten, H.A.B., 2018. Mushroom cultivation in the circular economy. Applied Microbiology and Biotechnology 102, 7795–7803. doi:10.1007/s00253-018-9226-8

Hackett, R., 2015. Spent mushroom compost as a nitrogen source for spring barley. Nutrient Cycling in Agroecosystems 102, 253–263. doi:10.1007/s10705-015-9696-3

Harrell, F., Dupont, C., 2016. Hmisc: Harrell Miscellaneous. R Package version 3.17-4.

Hellequin, E., Monard, C., Quaiser, A., Henriot, M., Klarzynski, O., Binet, F., 2018. Specific recruitment of soil bacteria and fungi decomposers following a biostimulant application increased crop residues mineralization. PLOS ONE 13, e0209089. doi:10.1371/journal.pone.0209089

Henry, R., Schang, C., Coutts, S., Kolotelo, P., Prosser, T., Crosbie, N., Grant, T., Cottam, D., O’Brien, P., Deletic, A., McCarthy, D., 2016. Into the deep: Evaluation of SourceTracker for assessment of faecal contamination of coastal waters. Water Research 93, 242–253. doi:10.1016/J.WATRES.2016.02.029

Jordan, S.N., Mullen, G.J., Murphy, M.C., 2008. Composition variability of spent mushroom compost in Ireland. Bioresource Technology 99, 411–418. doi:10.1016/j.biortech.2006.12.012

Kandeler, E., Gerber, H., 1988. Short-term assay of soil urease activity using colorimetric determination of ammonium. Biology and Fertility of Soils 6, 68–72. doi:10.1007/BF00257924

Keeney, D. R., and Nelson, D. W. (1982). Nitrogen—Inorganic forms. In “Methods of Soil Analysis— Part 2,” (A. L. Page, ed.), pp. 643–698. American Society of Agronomy, Madison, WI, USA.

Knights, D., Kuczynski, J., Charlson, E.S., Zaneveld, J., Mozer, M.C., Collman, R.G., Bushman, F.D., Knight, R., Kelley, S.T., 2011. Bayesian community-wide culture-independent microbial source tracking. Nature Methods 8, 761–763. doi:10.1038/nmeth.1650

Kurtz, Z.D., Müller, C.L., Miraldi, E.R., Littman, D.R., Blaser, M.J., Bonneau, R.A., 2015. Sparse and Compositionally Robust Inference of Microbial Ecological Networks. PLOS Computational Biology 11, e1004226. doi:10.1371/journal.pcbi.1004226

Lang, C., 1958. Simple microdetermination of Kjeldahl nitrogen in biological materials. Analytical Chemistry 30, 1692–1694. doi:10.1016/0026-265X(59)90212-7

Li, F., Chen, L., Zhang, J., Yin, J., Huang, S., 2017. Bacterial Community Structure after Long-term Organic and Inorganic Fertilization Reveals Important Associations between Soil Nutrients and Specific Taxa Involved in Nutrient Transformations. Frontiers in Microbiology 8, 187. doi:10.3389/fmicb.2017.00187

Ling, N., Zhu, C., Xue, C., Chen, H., Duan, Y., Peng, C., Guo, S., Shen, Q., 2016. Insight into how organic amendments can shape the soil microbiome in long-term field experiments as revealed by network analysis. Soil Biology and Biochemistry 99, 137–149. doi:10.1016/J.SOILBIO.2016.05.005

Lourenço, K.S., Suleiman, A.K.A., Pijl, A., van Veen, J.A., Cantarella, H., Kuramae, E.E., 2018. Resilience of the resident soil microbiome to organic and inorganic amendment disturbances and to temporary bacterial invasion. Microbiome 6, 142. doi:10.1186/s40168-018-0525-1

Martiny, J.B., Martiny, A.C., Weihe, C., Lu, Y., Berlemont, R., Brodie, E.L., Goulden, M.L., Treseder, K.K., Allison, S.D., 2017. Microbial legacies alter decomposition in response to simulated global change. The ISME Journal 11, 490–499. doi:10.1038/ismej.2016.122

Meyer, K.M., Petersen, I.A.B., Tobi, E., Korte, L., Bohannan, B.J.M., 2019. Use of RNA and DNA to Identify Mechanisms of Bacterial Community Homogenization. Frontiers in Microbiology 10, 2066. doi:10.3389/fmicb.2019.02066

Munson-McGee, J.H., Field, E.K., Bateson, M., Rooney, C., Stepanauskas, R., Young, M.J., 2015. Nanoarchaeota, Their Sulfolobales Host, and Nanoarchaeota Virus Distribution across Yellowstone National Park Hot Springs. Applied and Environmental Microbiology 81, 7860–7868. doi:10.1128/AEM.01539-15

Paul Chowdhury, S., Babin, D., Sandmann, M., Jacquiod, S., Sommermann, L., Sørensen, S.J., Fliessbach, A., Mäder, P., Geistlinger, J., Smalla, K., Rothballer, M., Grosch, R., 2019. Effect of long-term organic and mineral fertilization strategies on rhizosphere microbiota assemblage and performance of lettuce. Environmental Microbiology 21, 1462–2920.14631. doi:10.1111/1462-2920.14631

Paula, F.S., Tatti, E., Abram, F., Wilson, J., O’Flaherty, V., 2017. Stabilisation of spent mushroom substrate for application as a plant growth-promoting organic amendment. Journal of Environmental Management 196. doi:10.1016/j.jenvman.2017.03.038

Peay, K.G., von Sperber, C., Cardarelli, E., Toju, H., Francis, C.A., Chadwick, O.A., Vitousek, P.M., 2017. Convergence and contrast in the community structure of Bacteria, Fungi and Archaea along a tropical elevation–climate gradient. FEMS Microbiology Ecology 93. doi:10.1093/femsec/fix045

Philippot, L., Raaijmakers, J.M., Lemanceau, P., van der Putten, W.H., 2013. Going back to the roots: the microbial ecology of the rhizosphere. Nature Reviews Microbiology 11, 789–799. doi:10.1038/nrmicro3109

Randall, K., Brennan, F., Clipson, N., Creamer, R., Griffiths, B., Storey, S., Doyle, E., 2019. Soil bacterial community structure and functional responses across a long-term mineral phosphorus (Pi) fertilisation gradient differ in grazed and cut grasslands. Applied Soil Ecology 138, 134–143. doi:10.1016/J.APSOIL.2019.02.002

Roy, S., Barman, S., Chakraborty, U., Chakraborty, B., 2015. Evaluation of Spent Mushroom Substrate as biofertilizer for growth improvement of Capsicum annuum L. Journal of Applied Biology & Biotechnology 3, 22–27. doi:10.7324/JABB.2015.3305

Shade, A., Jones, S.E., Caporaso, J.G., Handelsman, J., Knight, R., Fierer, N., Gilbert, J.A., 2014. Conditionally rare taxa disproportionately contribute to temporal changes in microbial diversity. MBio 5, e01371–14. doi:10.1128/mBio.01371-14

Shand, C.A., Williams, B.L., Coutts, G., 2008. Determination of N-species in soil extracts using microplate techniques. Talanta 74, 648–654. doi:10.1016/j.talanta.2007.06.039

Strickland, M.S., Rousk, J., 2010. Considering fungal:bacterial dominance in soils e Methods, controls, and ecosystem implications. Soil Biology and Biochemistry 42, 1385–1395. doi:10.1016/j.soilbio.2010.05.007

Taylor, D.L., Walters, W.A., Lennon, N.J., Bochicchio, J., Krohn, A., Caporaso, J.G., Pennanen, T., 2016. Accurate Estimation of Fungal Diversity and Abundance through Improved Lineage-Specific Primers Optimized for Illumina Amplicon Sequencing. Applied and Environmental Microbiology 82, 7217–7226. doi:10.1128/AEM.02576-16

Varela-Cervero, S., López-García, Á., Barea, J.M., Azcón-Aguilar, C., 2016. Differences in the composition of arbuscular mycorrhizal fungal communities promoted by different propagule forms from a Mediterranean shrubland. Mycorrhiza 26, 489–496. doi:10.1007/s00572-016-0687-2

Wang, H., Wei, Z., Mei, L., Gu, J., Yin, S., Faust, K., Raes, J., Deng, Y., Wang, Y., Shen, Q., Yin, S., 2017. Combined use of network inference tools identifies ecologically meaningful bacterial associations in a paddy soil. Soil Biology and Biochemistry 105, 227–235. doi:10.1016/J.SOILBIO.2016.11.029

Wang, Y., Nagaoka, K., Hayatsu, M., Sakai, Y., Tago, K., Asakawa, S., Fujii, T., 2012. A novel method for RNA extraction from Andosols using casein and its application to amoA gene expression study in soil. Applied Microbiology and Biotechnology 96, 793–802. doi:10.1007/s00253-012-4342-3

Wei, T., Simko, V., 2017. R package “corrplot”: Visualization of a Correlation Matrix (Version 0.84). Available from https://github.com/taiyun/corrplot

White, T.J., Bruns, T., Lee, S., Taylor, J., 1990. AMPLIFICATION AND DIRECT SEQUENCING OF FUNGAL RIBOSOMAL RNA GENES FOR PHYLOGENETICS, in: PCR Protocols. Elsevier, pp. 315–322. doi:10.1016/B978-0-12-372180-8.50042-1

Wilhelm, R.C., Singh, R., Eltis, L.D., Mohn, W.W., 2019. Bacterial contributions to delignification and lignocellulose degradation in forest soils with metagenomic and quantitative stable isotope probing. The ISME Journal 13, 413–429. doi:10.1038/s41396-018-0279-6

Zhang, Y., Zhao, Z., Dai, M., Jiao, N., Herndl, G.J., 2014. Drivers shaping the diversity and biogeography of total and active bacterial communities in the South China Sea. Molecular Ecology 23, 2260–74. doi:10.1111/mec.12739

Zhou, J., Deng, Y., Luo, F., He, Z., Yang, Y., 2011. Phylogenetic molecular ecological network of soil microbial communities in response to elevated CO2. MBio 2, e00122–11. doi:10.1128/mBio.00122-11

